# Opposing Sequential Biases in Direction and Time Reproduction: Influences of Task Relevance and Working Memory

**DOI:** 10.1101/2024.01.28.577640

**Authors:** Si Cheng, Siyi Chen, Zhuanghua Shi

## Abstract

Our current perception and decision-making are shaped by recent experiences, a phenomenon known as serial dependence. While serial dependence is well-documented in visual perception and has been recently explored in time perception, their functional similarities across non-temporal and temporal domains remain elusive, particularly in relation to task relevance and working memory load. To address this, we designed a unified experimental paradigm using coherent motion stimuli to test both direction and time reproduction. The direction and time tasks were randomly mixed across trials. Additionally, we introduced pre-cue versus post-cue settings in separate experiments to manipulate working memory load during the encoding phase. We found attractive biases in time reproduction but repulsive biases in direction estimation. Notably, the temporal attraction was more pronounced when the preceding task was also time-related. In contrast, the direction repulsion remained unaffected by the nature of the preceding task. Additionally, both attractive and repulsive biases were enhanced by the post-cue compared to the pre-cue. Our findings suggest that opposing sequential effects in non-temporal and temporal domains may originate from different processing stages linked to sensory adaptation and post-perceptual processes involving working memory.

## Introduction

Our brain estimates the magnitude of stimuli not only from noisy sensory inputs but also from the recent past. This integration process results in sequential (serial) dependence and central tendency bias (Cicchini et al., 2023; Fischer & Whitney, 2014; Glasauer & Shi, 2022; Manassi et al., 2023; Pascucci et al., 2023). Sequential effects can lead to either attractive or repulsive biases (Ceylan & Pascucci, 2023; Cicchini et al., 2023; Czoschke et al., 2019; Manassi et al., 2023; Moon & Kwon, 2022), depending on the functional roles they serve. Sequential attraction occurs when consecutive stimuli are perceived as more similar than they actually are, believed to promote perceptual stability by integrating similar visual inputs over time (Cicchini et al., 2023; Fischer & Whitney, 2014; Glasauer & Shi, 2022; Liberman et al., 2016). However, there is a debate on whether it represents a purely perceptual phenomenon (Czoschke et al., 2019; Fornaciai & Park, 2018; Murai & Whitney, 2021; Pascucci et al., 2024) or relies more on memory traces of previous stimuli (Bae & Luck, 2020; Ceylan et al., 2021; Ceylan & Pascucci, 2023; Pascucci et al., 2019).

Sequential repulsion, on the other hand, occurs when the current perception is biased away from the preceding or concurrent stimulus. This effect usually appears when multiple stimuli need to be held in working memory (Czoschke et al., 2019) or when stimuli are presented but not reported (Ceylan & Pascucci, 2023; Pascucci et al., 2019; Pascucci & Plomp, 2021). It is thought to amplify small but potentially important differences between stimuli (Burr & Cicchini, 2014), maximizing discriminability (Czoschke et al., 2019; Fritsche et al., 2017) and perceptual accuracy (Ceylan & Pascucci, 2023; Fritsche et al., 2020; Moon & Kwon, 2022).

Sequential effects have been indicated to arise from perceptual or post-perceptual decision stages (Bae & Luck, 2020; Ceylan et al., 2021; Ceylan & Pascucci, 2023; Fischer et al., 2020; Fornaciai & Park, 2019; Fritsche & de Lange, 2019b; Pascucci et al., 2019). The perceptual perspective suggests that sequential attraction helps maintain perceptual stability and temporal continuity by integrating past and current information to filter out abrupt noises (Czoschke et al., 2019; Fornaciai & Park, 2018; Murai & Whitney, 2021; Pascucci et al., 2024). The evidence comes from the fact that this bias can even emerge in the absence of a decision process when participants are not required to report the target feature in the previous trial (Czoschke et al., 2019; Fornaciai & Park, 2018; Murai & Whitney, 2021). Post-perceptual perspective links sequential effects to decision-related factors (Bae & Luck, 2020; Ceylan et al., 2021; Ceylan & Pascucci, 2023; Fischer et al., 2020; Fornaciai & Park, 2019; Fritsche & de Lange, 2019b; Pascucci et al., 2019). For example, in a study involving orientation or color judgments with a post-cue, sequential dependence emerged only when both the preceding and current trials were the same task, diminishing when tasks differed (Bae & Luck, 2020). These findings suggest that task-related responses from previous trials, not just the encoding of the prior stimulus, are necessary for sequential effects.

The two opposing effects, attraction and repulsion, have historically been studied separately, recent research suggests they may co-occur and interact during perceptual processing (Feigin et al., 2021; Fritsche et al., 2017; Moon & Kwon, 2022; Pascucci et al., 2019; Pascucci & Plomp, 2021; Sadil et al., 2024; Sheehan & Serences, 2023; Zhou et al., 2024). For instance, a recent study on motion direction showed that the preceding direction response induced an attractive bias, while the preceding motion direction caused a repulsion bias, both contributing to serial dependence (Moon & Kwon, 2022). The previous response, being the observer’s final estimate of the prior stimulus, serves as a predictor for upcoming stimuli, influencing the current perceptual estimate (Burr & Cicchini, 2014; Moon & Kwon, 2022; Sadil et al., 2024). Thus, attraction could also operate at the decision response stage or higher-level processes. Conversely, adaptation in the neural population responsible for encoding specific features, such as orientation or motion direction, plays a crucial role in repulsion (Alais et al., 2017; Fritsche et al., 2017, 2020; Moon & Kwon, 2022; Sheehan & Serences, 2022). This adaptation, often manifested as a reduction in neural response gain to the previously encoded feature, leads to a repulsive shift in the perceived stimulus away from the preceding one or the long-term past (Fritsche et al., 2017, 2020; Moon & Kwon, 2022).

The role of working memory in serial dependence is crucial and multifaceted, as it influences decision-making and motor planning (Bae & Luck, 2020; Bliss et al., 2017; de Azevedo Neto & Bartels, 2021; Kiyonaga, Scimeca, et al., 2017). Research shows that increasing temporal delay between stimulus presentation and response, thereby imposing greater working memory loads, leads to a stronger attractive bias toward the preceding stimulus (Bliss et al., 2017). A recent fMRI study on duration reproduction further demonstrates that consecutive active responses, compared to passive viewing, enhanced attractive biases (Cheng et al., 2023). Crucially, in these consecutive response trials, sequential effects negatively correlated with hippocampus activities (Cheng et al., 2023). While increased working memory demands generally amplify attractive biases, evidence suggests a more complex relationship with repulsive biases. Some findings indicated that repulsive biases might arise from sensory adaptation occurring at an early stage of visual processing, potentially independent of working memory (Fritsche et al., 2017; Kiyonaga, Scimeca, et al., 2017).

It is important to note that the aforementioned studies on sequential effects have predominantly concentrated on non-temporal features within the visual domain, where low-level sensory adaptation and higher-level memory processing may interact. Research on sequential effects in time perception is less developed, with main findings often suggesting attractive rather than repulsive biases (Glasauer & Shi, 2022; Wehrman et al., 2018, 2023; Wiener et al., 2014). The prevalence of attractive biases in time perception remains unclear, but several distinct features in time perception might contribute to this predominance. In contrast to visual processing, time perception is not bound to a specific sensory system (Wittmann & Paulus, 2008) and may rely more heavily on the representation of stimuli in working memory (B.-M. Gu et al., 2015; Shi et al., 2013; Teki & Griffiths, 2016). Sensory adaptation-induced repulsion biases might be specific to certain modalities, such as vision. Additionally, unlike some visual features that can be simultaneously presented and held in memory, time intervals are monitored and processed sequentially. The natural stimulus distribution also differs between non-temporal visual features (e.g., orientation, motion direction, color) and magnitude stimuli (e.g., time, numerosity, length) (Hahn & Wei, 2024). For example, motion direction follows a circular distribution, ranging from 0 to 360 degrees, potentially mitigating the influence of central tendency biases, whereas temporal stimuli follow an open-scale distribution, mainly influenced by central tendency biases and sequential (serial) dependence biases (Glasauer & Shi, 2022).

Given these distinct processes for non-temporal visual features and time, it remains uncertain if sequential effects share the same mechanisms across temporal and non-temporal domains. Notably, working memory is often shared between tasks across domains, so its influence on sequential effects might have some commonalities. However, the contribution of working memory to sequential effects may differ for temporal and non-temporal processes, which remains an open issue. Studies concurrently addressing sequential effects in both spatial features (such as motion direction) and time are particularly rare. This study aims to address this gap by investigating sequential effects in both temporal and non-temporal domains using coherent motion stimuli, involving both direction and time reproduction tasks.

We hypothesized that sensory adaptation is specific to sensory systems, so adaptation-induced repulsion bias is likely to occur in the non-temporal rather than temporal domain. Early studies suggest that sensory adaptation and perceptual level serial dependence require minimal working memory (Fischer & Whitney, 2014; Fritsche et al., 2017), indicating that sequential effects that emerge at early perceptual processing may be resistant to task changes. In contrast, temporal processing heavily involves memory processing, and its sequential effects may be enhanced by task relevancy and working memory loads. To test these hypotheses, we designed two experiments employing the pre-cue vs. post-cue setting, the latter used in prior research (Bae & Luck, 2020; Cheng et al., 2023). Participants reproduced either the direction (direction task) or duration (time task) of a coherent motion display based on a cue shown either before (pre-cue, Experiment 1) or after (post-cue, Experiment 2) stimulus presentation. By using this unified experimental paradigm involving both direction and time reproduction tasks, we aimed to compare the non-temporal and temporal sequential effects. By randomly interleaving the direction and time tasks, we expected to observe task-relevant modulation for post-perceptual serial dependence. Additionally, comparing the pre-cue and post-cue results, we anticipated an enhancement of sequential effects by memory load.

To foreshadow the results, in two experiments (each N=23), we found that the post-cue, relative to the pre-cue, enhanced sequential effects. For timing tasks, strong attraction was observed only when the task of the preceding trial was the same, indicating that task consistency enhanced the attraction bias. In contrast, we observed repulsion biases for mild inter-trial orientation differences, regardless of the preceding task. These findings highlight the differential mechanisms underlying working memory-based temporal assimilation and adaptation-based spatial repulsion in direction, both of which are reinforced by working memory.

## Experiment 1

### Method

#### Participant

Twenty-three volunteers participated in Experiment 1 (14 females, 9 males; age ranged from 21 to 38, mean ± SD: 26.83 ± 4.18 years). All participants were right-handed and had normal or corrected-to-normal visual acuity and color vision. A meta-analysis of serial dependence in orientation reported in 35 different publications revealed a substantial effect size with a media Fisher *z_r_* of 0.66 (equivalent to Cohen’s d of 1.416) (Manassi et al., 2023). The difference in serial dependence between post-cue conditions (action vs. no action) in a recent duration reproduction task with a similar setup (Cheng et al., 2023) yields a large difference (Cohen’s *d* = *1.367*). To determine the appropriate sample size for our study, we conducted a priori power analysis using G*Power 3 (Faul et al., 2007). Considering our experimental design, which included a pre-cue versus post-cue manipulation and various tasks, we adopted a conservative approach. Based on half of the reported effect size (*d* = 0.7) and a significance level of α = .05, we aimed for a statistical power of 80% (1-β). Our calculations revealed that a minimum of 15 participants would be required to achieve the desired level of statistical power. On the safe side, we increased the sample size to 23. Prior to the experiment, participants gave written informed consent and received compensation of 9 Euro/hour for their involvement. The study was approved by the ethics committees of the Psychology Department at LMU Munich.

### Stimuli and procedure

We used PsychoPy3 (Peirce et al., 2019) to control stimulus presentation and data collection. Participants were seated in a soundproof, dimly lit cabin, resting their heads on a chin rest. They kept a viewing distance of 60 cm from a 21-inch CRT monitor (refresh rate at 85 Hz), which presented stimuli on a light grey display background (luminance of 39.3 cd/m^2^).

Figure 1 illustrates the study setup. We employed a pre-cue for the time and motion direction reproduction task in Experiment 1. Each trial began with a fixation dot (subtended 0.5°; luminance of 85.7 cd/m^2^) for 500 ms, indicating the beginning of the trial and prompting participants to focus their attention. A pre-cue with a letter (“D” or “T”, visual angle of 0.8° × 1.0°, luminance of 85.7 cd/m^2^) appeared in the display center for 500 ms, indicating whether participants should report the direction (the letter “D”) or time (the letter “T”) of the coherent motion. The reproduction task consisted of an encoding phase and a reproduction phase. Immediately after the cue display, the encoding phase started with a random dot kinematogram (RDK) display for 400 to 600 ms. The RDK consisted of 100 randomly generated white dots (each dot diameter of 0.2°; luminance of 85.7 cd/m^2^) within a dark disk (subtended 17.8°, luminance of 16.5 cd/m^2^) at the center of the screen. Those random dots randomly walked at a speed of 1°/s. Subsequently, the white dots turned green (luminance of 45.8 cd/m^2^) and moved coherently (100%) with a speed of 6°/s in one direction (randomly selected from 11.25° to 348.75°, in steps of 22.5°) for a randomly sampled duration (0.8, 1.0, 1.2, 1.4, or 1.6 s). Since the primary objective of our research was to investigate the sequential effects in motion direction and time tasks, we selected this stimulus duration range to keep underlying mechanisms of time perception stable (e.g., sub- and supra-second time perception are often assumed to rely on different mechanisms and different brain regions (Hayashi et al., 2014; Lewis & Miall, 2003) as well as balance task difficulty among trials. Additionally, the cardinal motion directions (0°, 90°, 180°, 270°) were also avoided to rule out the cardinal rules (the orientation judgments were more accurate at horizontal and vertical orientation (Bae, 2024; Girshick et al., 2011; Mao & Stocker, 2021).

**Figure 1.**
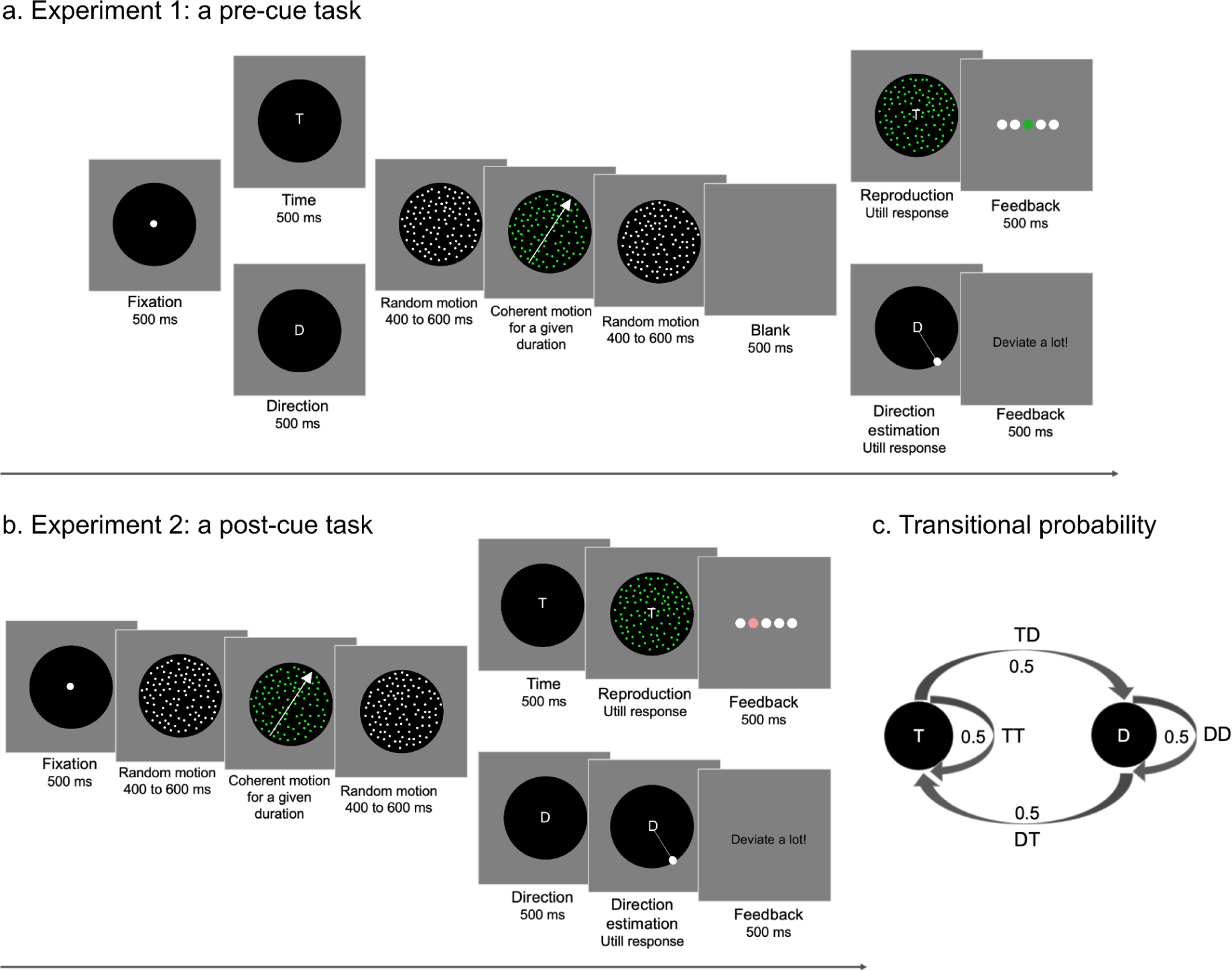
Schematic illustration of the study. (**a**) Procedure of Experiment 1 (the pre-cue task). Each trial began with a fixation dot, followed by a pre-cue letter indicating whether participants should report the direction (“D”) or time (“T”) of motion. The encoding phase began with a random dot kinematogram (RDK). When the white random-walked dots turned green, they moved in a specific direction for a given duration (indicated by the illustrative white arrow, not shown in the experiment). Then, they reverted to the random-walked white dots motion. The coherent motion was the target stimulus that participants had to remember either its direction or time according to the cue. The reproduction phase began after 500 ms of a blank screen. For the time task, a green RDK display appeared, and participants had to click the left mouse to terminate it when the duration matched to the coherent movement perceived during the encoding phase. For the direction task, a display with an adjustable line point appeared, and participants adjusted its orientation using a mouse, finalizing the direction report by pressing the space key. After their response, a 500-ms feedback display showed their accuracy; (**b**) Procedure of Experiment 2 (the post-cue task). It included the same encoding phase as in Experiment 1 but without a pre-cue. After the encoding phase, a post-cue displays for 500 ms, prompting participants for either time or direction reproduction. The rest of the procedure was the same as in Experiment 1. (**c**) The inter-trial transitional probability (from trial *n-1* to trial *n*) between the time and direction tasks. This transitional structure guarantees equal transitional probability of “Direction to Time (DT)”, “Time to Time (TT)”, “Direction to Direction (DD)”, and “Time to Direction (TD)” trials.

When a dot moved out of the dark disk, a new dot was randomly regenerated within the disk, keeping 100 dots all the time for the RDK presentation. This coherent motion was the encoding target. According to the pre-cue letter, participants were instructed to remember the direction or duration of the coherent motion for their reproduction task. Then, the dots reverted to random-walked white dot movements for 400 to 600 ms. The RDK displays served as masks before and after the target.

Following the encoding phase, there was a 500 ms blank interval before the participants’ response. For the time task, the letter ‘T’ appeared in the center of the screen along with random-walked green dots (100 dots, each dot diameter of 0.2°; luminance of 45.8 cd/m^2^; velocity of 1 °/s). We used a random-walked motion instead of a coherent motion to minimize inter-trial bias caused by the motion from the reproduction phase, given that we were interested in the sequential effects of the motion from the encoding phase. Participants had to terminate the presentation by clicking the left mouse when they perceived the duration to match the duration of the green dots’ coherent motion from the encoding phase. After their response, feedback was given for 500 ms using a horizontally arranged display of five disks (each subtended 1.8°). The accuracy of the response was indicated by one colored disk, from the left to the right, indicating the relative error below −30%, between [-30%,-5%], (−5%, 5%), [5%, 30%] and greater than 30%, respectively. The middle circle appeared in green, representing high accuracy. The middle left and middle right circles are in light red, indicating some deviation from the actual interval. The utmost left and right circles were shown in dark red, indicating a large reproduction error. If it was a direction task, a line pointer with a superimposed letter D appeared. The line started from the center, pointing in a random direction. Participants adjusted the pointer by moving the mouse and confirmed the final orientation by pressing the space key. If the final estimated direction deviated more than 60° from the true direction, a feedback display with the message “Direction deviated a lot!” appeared at the center of the display for 500 ms. Otherwise, a blank display appeared for 500 ms. The next trial began after a second intertrial interval.

Before the formal experiment, participants received 16 practice trials to familiarize themselves with the task. The formal experiment consisted of 480 trials, randomly shuffled with half for time reproduction and half for direction reproduction. The inter-trial transitional probability (from trial *n-1* to trial *n*) between the time and direction trials ensured an equal probability of “Direction to Time (DT)”, “Time to Time (TT)”, “Direction to Direction (DD)”, and “Time to Direction (TD)” trials. Participants could take a short break after each block of 30 trials.

### Data analysis

We excluded the first trial of each block, resulting in 16 omitted trials. Outliers due to accidental button presses or inattention were also excluded, specifically those with reproduction errors exceeding three standard deviations from the mean error for time report trials and response errors larger than 45° for direction report trials, before proceeding with further analyses. These outliers were rare, constituting only 0.38% of time reproduction trials (ranging individually from 0 to 6 outlier trials) and 0.24% of direction report trials (ranging individually from 0 to 13 outlier trials). Next, we categorized the remaining trials into four categories based on the inter-trial transition (from trial *n-1* to trial *n*): Direction to Time (DT), Time to Time (TT), Direction to Direction (DD), and Time to Direction (TD). To investigate the influence of prior tasks on the sequential effects of current estimates, we conducted separate analyses for time reproduction and direction reproduction trials.

#### Time reproduction trials

We focused on two conditions: the “Time to Time (TT)” representing the prior task-related condition and the “Direction to Time (DT)” representing the prior task-unrelated condition. Previous research has shown that subjective timing, on an open scale, is susceptible to both the central tendency bias and the sequential bias (Glasauer & Shi, 2022; Holland & Lockhead, 1968). The central tendency bias leads to an overestimation of shorter durations and an underestimation of longer durations, while the sequential bias indicates that duration estimations are influenced by preceding durations.

To calculate the central tendency effect, we employed linear regression to approximate the relationship between the current reproduction error (*Error_n_*) and the current duration (*T_n_*) (Cicchini et al., 2012; Jazayeri & Shadlen, 2010; Shi et al., 2013).

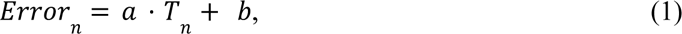

where the absolute slope of the linear fit (|*a|*) reflects the central tendency effect. A slope of 0 indicates no central tendency, and 1 represents a strong central tendency.

The conventional measures of the serial dependence effect, which correlates the current response error to the difference between the previous and the current stimuli (Bliss et al., 2017; Cicchini et al., 2018; e.g., Fischer & Whitney, 2014; Kiyonaga, Manassi, et al., 2017), are not sufficient to separate sequential dependence from the central tendency bias (for more details, see Glasauer & Shi, 2022). Thus, we employed linear regression to the previous trial (Holland & Lockhead, 1968; Jesteadt et al., 1977) to analyze the sequential effect. This involved examining the correlation between the current error and the previous duration (*T_n_*_-1_):

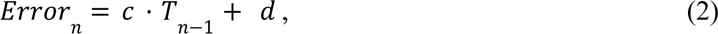

where the slope of the linear fit (*c*) indicates the sequential bias (e.g., Glasauer & Shi, 2022; Jesteadt et al., 1977). A positive slope indicates that the current estimation is attracted towards the previous duration (also called the “assimilation”). In contrast, a negative slope indicates that the current time estimation is repulsed from the previous duration. Additionally, as a sanity check and for further verification, we also computed the sequential effect using similar regressions for the durations presented in future trials (n+1).

#### Direction estimation trials

We focused on two conditions: the “Direction to Direction (DD)” representing the prior task-related condition and the “Time to Direction (TD)” representing the prior task-unrelated condition. The direction of motion was randomly selected from a circular distribution featuring 16 equally spaced angles (from 11.25° to 348.75°, in steps of 22.5°), effectively neutralizing any central tendency. Consequently, we focused solely on the sequential effect and skipped the central tendency analysis. The response error was calculated as the difference between the reported direction and the true motion direction for the current trial (i.e., estimate - direction). Negative errors indicated a counter-clockwise deviation from the true direction, while positive errors suggested a clockwise deviation. Additionally, the direction difference was also calculated between the current trial and the previous trial (the previous direction - the current direction), following the same method used in prior research (e.g., Fischer & Whitney, 2014). Trials with a direction difference of 0° or ± 180° were excluded, as response errors relative to these direction differences are undefined. Following previous research (Moon et al., 2022) highlighting a significant role of non-directional orientation in the coding of visual motion direction, we reduced the direction difference range from [-180 to 180°] to [-90 to 90°] accordingly. To better reflect the repulsion and attractive biases, we converted the response errors from clockwise or counterclockwise directions to the repulsion (negative) and attractive (positive) biases by collapsing the direction differences to the positive range [0, 90.0°]. This analysis resembles the analysis of previous studies (Bae & Luck, 2020).

Prior research has shown that small orientation differences (within 45°) led to a significant attractive bias, while large orientation differences (greater than 45°) resulted in a repulsive bias (Bae & Luck, 2019; Bliss et al., 2017; Fritsche et al., 2017; Fritsche & de Lange, 2019b; Samaha et al., 2019). For example, attraction was observed when the orientation difference was around 17°, and repulsion was found for orientations more than 60° apart (Fritsche et al., 2017). We hypothesized that the reported orientation for the current trial would be repulsed by the orientation in the previous trial, particularly when orientations differed markedly. Thus, we calculated the average response errors for mild-orientation-difference trials (45.0° and 67.5°) and compared them to zero for each condition (the prior task being direction reproduction or time reproduction for the current direction reproduction trials).

Additionally, we computed the sequential effect in direction estimation, separated for the “short” and “long” stimulus presentation. Durations of 0.8 and 1.0 seconds were categorized as “short”, while durations of 1.4 and 1.6 seconds were deemed “long”. We excluded the intermediate duration of 1.2 seconds in this analysis. We then analyzed whether the sequential effects in direction estimation were modulated by the durations (short vs. long) of the target stimulus.

The statistical significance of the central tendency effect and the sequential effect was assessed individually using analysis of variances (ANOVAs) and one-sample *t*-tests against a null hypothesis of zero effect. Paired *t*-tests were run for within-subject between-condition comparisons.

## Results and discussion

### Time Reproduction

Overall, the mean reproduction errors (and associated standard errors, SEs) for prior time reproduction trials (task-related: TT) and prior direction estimation trials (task-unrelated: DT) were 51 ± 14 ms and 57 ± 15 ms, respectively. There was no significant difference between the two conditions (*t_(22)_* = −0.647, *p* = .524, *d* = −0.090). To examine the precision of duration reproduction for two kinds of preceding task (time vs. direction), we calculated the standard deviation of reproduction between TT and DT conditions, and it didn’t show any significant difference between the two conditions (*t_(22)_* = 0.142, *p* = .888, *d* = 0.012). The central tendency effect (Eq. 1) was evident in both the prior task-related TT and prior task-unrelated DT conditions. The mean central tendency indices (and associated SEs) were 0.317 ± 0.037 (*t*_(22)_ = 8.605, *p* <.001, *d* = 1.794) and 0.297 ± 0.039 (*t*_(22)_ = 7.565, *p* <.001, *d* = 1.577) for the TT and DT conditions, respectively. However, there was no difference between the conditions (*t*_(22)_ = 1.364, *p* = .186, *d* = 0.109, *BF*_10_ = 0.479), as illustrated in Figure 2a.

**Figure 2.**
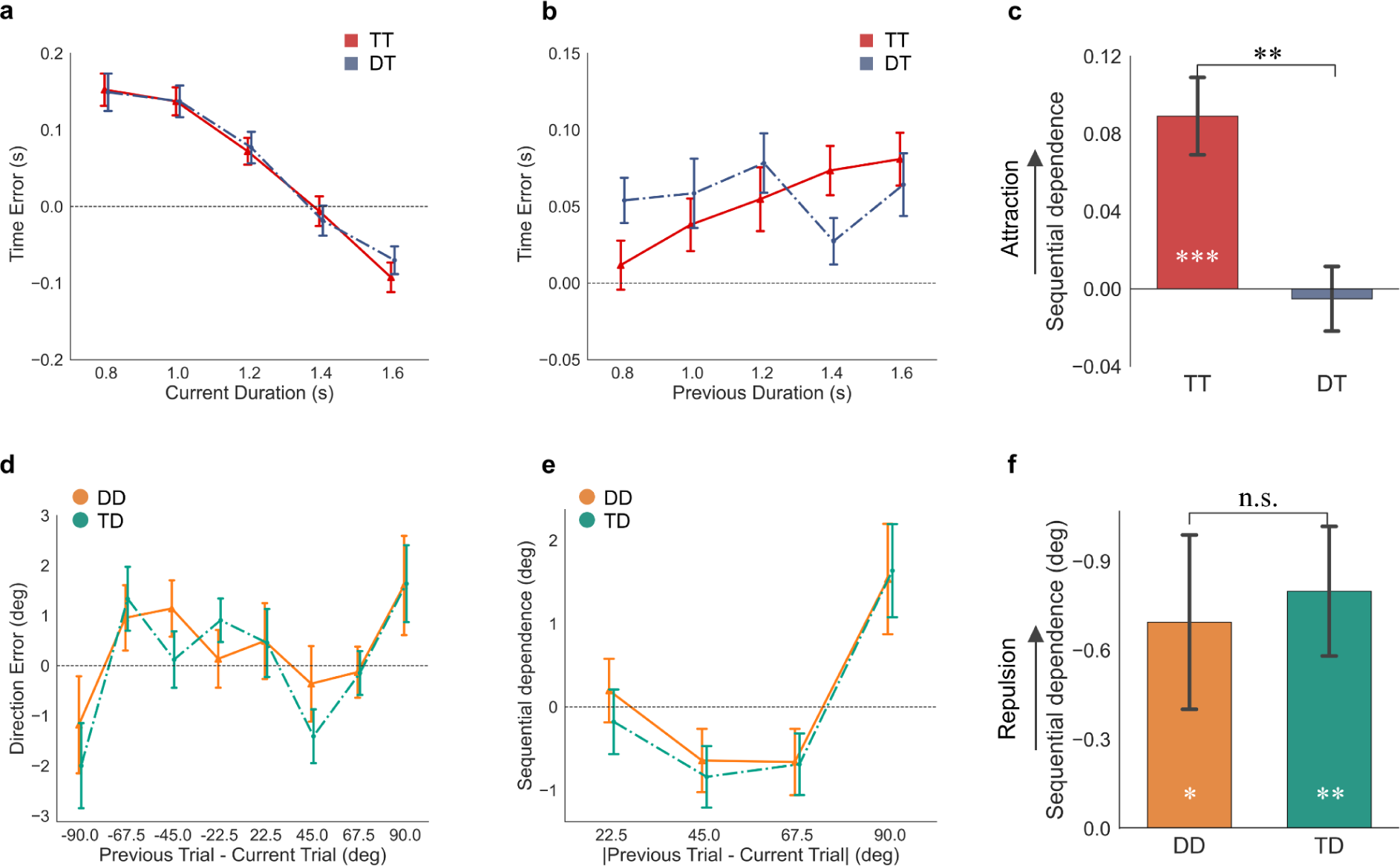
Results of Experiment 1. (**a**), (**b**), and (**c**) were the results of time reproduction trials. (**a**) Central tendency effect. Mean reproduction errors on the current sample duration are plotted separately for trials preceded by time report (TT) and direction report (DT). (**b**) Sequential effect. Mean reproduction errors on the previous duration, plotted separately for TT and DT conditions. (**c**) The mean slope of the linear fit. (**d**), (**e**), and (**f**) were the results of direction reproduction trials. (**d**) Mean response errors on the orientation difference of [-90°, 90°], plotted separately for trials preceded by direction report (DD) and time report (TD). The angular difference was realigned to represent the relative motion orientation (plus 180° for the opposite direction) of the previous trial. (**e**) Mean errors on the absolute orientation difference of [0°, 90°], plotted separately for DD and TD conditions. The sign of the response error was coded so that positive values indicate that the current-trial direction report was biased toward the direction of the previous trial, and negative values indicate that the current-trial direction report was biased away from the direction of the previous trial. Maximum repulsion occurred at 45° and 67.5° orientation differences. (**f**) Mean errors averaged across 45.0° and 67.5° were plotted separately for DD and TD conditions. Error bars represent ± SEM. ** denotes *p* < .01,* *p* < .05, and n.s. non-significant.

To quantify the sequential effect, we plotted the reproduction errors on the previous durations separately for the TT and DT conditions (as shown in Figure 2b). The reproduction error increased with increasing prior duration, showing that a longer prior duration attracted bias to a positive direction, manifesting an attractive sequential effect. This effect was quantified using the slope of linear regression (Eq. 2), which showed that the average slope was larger for the prior task-related TT condition (8.9% ± 2.0%) than that for the task-unrelated DT condition (−0.5% ± 1.7%), with paired t-test *t*_(22)_ = 3.813, *p* = .001, *d* = 1.064. The slope was only significantly positive for the TT condition (*t*_(22)_ = 4.457, *p* < .001, *d* = 0.929), but not for the DT condition (*t*_(22)_ = −0.304, *p* = .764, *d* = 0.063), as illustrated in Figure 2c. To ensure the validity of the findings and avoid statistical artifacts (Cicchini et al., 2014), we also tested and found no sequential effect on the durations presented in future trials (*ps* > .417).

### Direction Estimation

The mean response errors were plotted against the direction difference between the previous and the current trials (ranging from −90° to 90°, a positive value representing the difference in the clockwise direction), separated for the prior task-related (DD) and task-unrelated (TD) conditions (Figure 2d). We first examined the precision of response (the standard deviation of direction reproduction) between DD and TD conditions didn’t show any significant difference between the two conditions (*t_(22)_* = 0.967, *p* = .344, *d* = 0.061). Additionally, the direction errors were translated into the repulsion (negative) and attractive (positive) sequential effect and replotted as a function of the absolute orientation difference for each condition (illustrated in Figure 2e). By visual inspection, the maximum repulsion effect is likely between 45.0° and 67.5°, with a large difference between preceding trial types. Indeed, the average repulsive biases across the orientation differences of 45.0° and 67.5° were both significant negative (DD: −0.695° ± 0.294°, *t_(22)_* = 2.367, *p* =.027, *d* = 0.494; TD: −0.799° ± 0.218°, *t_(22)_* = 3.665, *p* =.001, *d* = 0.764), but no difference between the two, *t*_(22)_ = 0.283, *p* =.78, *d* = 0.084, *BF*_10_ = 0.227 (Figure 2f). To ensure the validity of the findings and avoid statistical artifacts (Cicchini et al., 2014), we also tested and found no repulsion effect across the orientation differences of 45.0° and 67.5° between the future (*n+1*) and current trials (*ps* > .635).

The sequential effect at 90° were significantly positive in both DD (1.537° ± 0.663°, *t_(22)_* = 2.320, *p* =.030, *d* = 0.484) and TD (1.638° ± 0.599°, *t*_(22)_ = 2.927, *p* =.008, *d* = 0.610) trials, but comparable between the two (*t*_(22)_ = −0.100, *p* =.921, *d* = −0.034, *BF*_10_ = 0.220). This condition at 90° was a special scenario. If participants’ judgments consider only the orientation, the effect can be interpreted as either attraction or repulsion. However, if judgments include both the orientation and direction), the effect is an attraction, assimilating toward the preceding direction. Previous research (Moon et al., 2022) indicated the significant role of non-directional orientation in the coding of visual motion direction. Thus, when the difference between the initial and the subsequent motion directions is 90°, attraction to the initial motion direction may be perceived as repulsion to the opposite direction if the motion direction is encoded in a non-directional orientation framework. The sanity check with the attraction effect for the orientation difference at 90° between the future (*n+1*) and current trials revealed no effects (*ps* > .372).

To rule out potential impacts of the presentation durations on sequential effects (shown at the direction differences: 45.0°, 67.5°, and 90°), we conducted three-way repeated measures ANOVA on response biases, considering current Stimulus Exposure^1^, Prior Task, and inter-trial Direction Difference (45.0°, 67.5°, and 90°) as main factors. The analysis did not reveal any significant effects of Stimulus Exposure or its related interactions 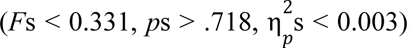. This suggests that variations in stimulus durations in our setup (800 - 1600 ms) did not affect the sequential effect.

Thus, Experiment 1 revealed an attractive bias in time reproduction only when the prior task was also time-related, suggesting that assimilation in temporal perception requires the previous trial to involve the same task. In contrast, the direction task yielded comparable biases unaffected by the prior task type and stimulus exposure. When the difference between two consecutive orientations was large (e.g., 45.0° and 67.5°), a repulsion bias was observed. For orientation differences (90°), an attractive bias emerged. The attractive effects at a 90° orthogonal difference might also be interpreted as a repulsion to the opposite motion direction.

### Experiment 2

In Experiment 1, we employed a pre-cue setting in which participants were aware of the response dimension in advance, and the unattended dimension did not require active working memory maintenance during encoding. This setup might have led to a diminished encoding of the unattended feature dimension, leading to the lack of sequential dependence. To address this, in Experiment 2, we adopted a post-cue setting where both response dimensions, time and direction, had to be memorized during the encoding phase, and a post-cue revealed which dimension was relevant for the response.

## Methods

### Participant

Twenty-three participants were recruited in Experiment 2 (13 females, 10 males; age 19 - 40, mean ± SD: 27.78 ± 5.31 years). All participants provided their written informed consent prior to the experiment and received 9 Euro/hour for their participation. The study was approved by the ethics committees of the Psychology Department at LMU Munich.

### Stimuli and procedure

Experiment 2 used the post-cue setting, which was essentially the same as in Experiment 1, with one key difference: the task cue display (500 ms) was shown after the encoding phase (see Figure 1 b). In order to perform the tasks properly, participants had to remember both direction and time in the encoding phase and then report one of them according to the post-cue.

### Data analysis

The data analysis for Experiment 2 followed essentially the same approach as that of Experiment 1. The first trial of each block was excluded. The outliers, using the same criteria as in Experiment 1, were rare in Experiment 2, on average only 0.40% of the time reproduction trials (ranging individually from 0 to 4 outlier trials) and 0.96% of the direction reproduction trials (ranging individually from 0 to 37 outlier trials).

## Results and discussion

### Time Reproduction

The average reproduction errors and their associated SEs for trials with prior time reproduction (task-related: TT) and trials with prior direction reproduction (task-unrelated: DT) were 34 ± 18 ms and 51 ± 16 ms, respectively. The difference between the two conditions was not significant, (*t*_(22)_ = −1.755, *p* = .093, *d* = −0.208, *BF*_10_ = 0.818). The standard deviation of duration reproduction between TT and DT conditions didn’t show any significant difference between the two conditions (*t_(22)_* = 0.316, *p* = .755, *d* = 0.020). Both conditions exhibited significant central tendency biases: 0.471 ± 0.057 and 0.486 ± 0.058 for the TT and DT conditions, respectively (*t*s_(22)_ > 8.3, *p*s <.001, *d*s > 1.7). There was no significant difference between them (*t*_(22)_ = −0.581, *p* = .580, *BF_10_* = 0.255, Figure 3a).

**Figure 3.**
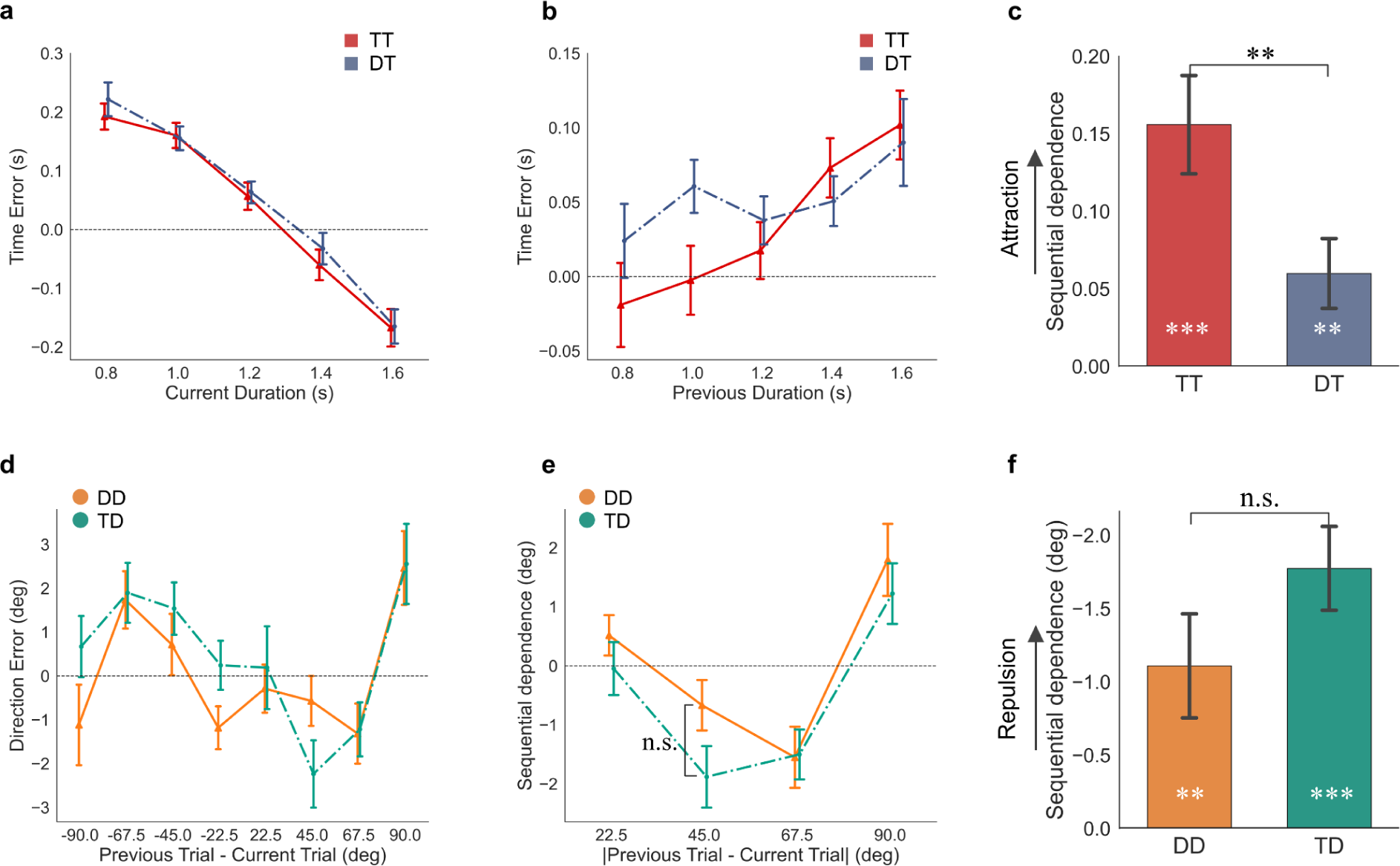
Results of Experiment 2. (**a**), (**b**), and (**c**) were the results of time reproduction trials. (**a**) Central tendency effect. Mean reproduction errors on the current sample duration, plotted separately for trials preceded by time report (TT) and direction report (DT). (**b**) Sequential effect. Mean reproduction errors on the previous duration, plotted separately for TT and DT conditions. (**c**) The slope of the linear fit. (**d**), (**e**), and (**f**) were the results of direction reproduction trials. (**d**) Mean response errors on the orientation difference of [-90°, 90°], plotted separately for trials preceded by direction report (DD) and time report (TD). The angular difference was realigned to represent the relative motion orientation (plus 180° for the opposite direction), rather than the motion direction, of the previous trial. (**e**) Mean errors on the absolute orientation difference of [0°, 90°], plotted separately for DD and TD conditions. The sign of the response error was coded so that positive values indicate that the current-trial direction report was biased toward the direction of the previous trial, and negative values indicate that the current-trial direction report was biased away from the direction of the previous trial. The difference between the DD and TD conditions for the 45° orientation difference didn’t reach significance, for the large separation was due to one participant who had large response errors in this type. Maximum repulsion occurred at 45° and 67.5° orientation differences. (**f**) Mean errors averaged across 45.0° and 67.5°, were plotted separately for DD and TD conditions. Error bars represent ± SEM. ** denotes *p* < .01, n.s no significant.

The sequential effects for each condition were illustrated in Figure 3b, with mean slopes of 15.6% ± 3.2% and 6.0% ± 2.3% for the task-related (TT) and task-unrelated (DT) conditions, respectively (Figure 3c). Both slopes were significantly higher than zero (*t*s_(22)_ > 2.6, *p*s < .02, *d*s > 0.55), indicating a significant attractive bias in both conditions. Moreover, a paired *t*-test showed that the assimilation was significantly larger in the TT than in the DT condition (*t*_(22)_ = 2.513, *p* = .020, *d* = 0.728). Tests of the sequential effect on future trial durations (*n+1*) ruled out any statistical artifacts (*ps* > .769).

### Direction Estimation

Figure 3d depicted the response errors against the direction difference from −90° to 90° for prior direction reproduction and time reproduction trials separately. The standard deviation of direction reproduction between prior direction report and time report trials didn’t show any significant difference between the two conditions (*t_(22)_*= 1.091, *p* = .287, *d* = 0.103). The direction errors were translated into the repulsion (negative) and attractive (positive) sequential effect and replotted in Figure 3e. Repulsion biases occurred at orientation differences of 45.0° and 67.5°. The averaged repulsion biases from the two orientation differences were significant for both task-related (DD) and task-unrelated (TD) conditions (DD: −1.110° ± 0.355°; TD: −1.775° ± 0.286°, *t*s*_(22)_* > 3.127, *p*s < .005, *d*s > 0.65), but did not differ from each other (*t_(22)_* = 1.918, *p* =0.068, *d* = 0.431, *BF_10_* = 1.038), as illustrated in Figure 3f.

We also observed an attractive bias for both TD and DD conditions at an orientation difference of 90° (DD: 1.796° ± 0.610° and TD: 1.224° ± 0.514°, *t*s*_(22)_* > 2.38, *p* < .026, *d* > 0.49), and they were comparable (*t_(22)_* = 0.937, *p* =.359, *d* = 0.211, *BF_10_* = 0.324). For sanity check, we tested the repulsion effect (at orientation differences of 45.0° and 67.5°) and attraction effect (at an orientation difference of 90°) across differences between the future (*n+1*) and current trials, and we found no effects (*ps* > .151).

Similar to Experiment 1, we conducted a further three-way repeated measures ANOVA on the reproduction biases, considering factors of Stimulus Exposure (Short vs. Long), Prior Task (Direction vs. Time), and inter-trial Direction Difference (45.0°, 67.5°, and 90°) failed to reveal any significance 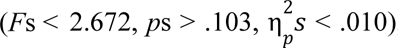, indicating the sequential bias was unaffected by the variations of stimulus exposure we used (800 to 1600 ms).

### Omnibus analysis

To gain a better understanding of the differences between the pre-cue and post-cue settings in terms of central tendency and sequential effects, we further conducted an omnibus analysis across both experiments. Specifically, we performed a two-way mixed ANOVA, with Prior Task (related task vs. unrelated task) as a within-subject factor and Cue Setting (Exp. 1: pre-cueing vs. Exp. 2: post-cueing) as a between-subject factor, on each effect of interest.

For the time reproduction trials, the two-way mixed measurement ANOVA on the central tendency index revealed a significant main effect of Cue Setting 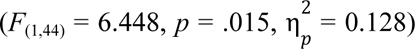, indicating a larger central tendency with the post-cue (Figure 4a). However, neither the main factor of Prior Task 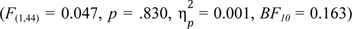 nor their interaction 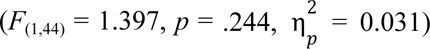 was significant. In contrast, the two-way mixed ANOVA on the sequential effect revealed both main factors were significant: Cue Setting, 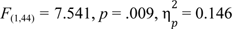; Prior Task, 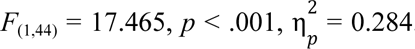. The results revealed that the attraction was significantly larger with the preceding same rather than the different task. Additionally, the attraction was amplified with the post-cue compared to the pre-cue (Figure 4b). However, there was no significant interaction 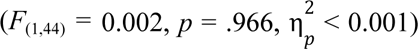. Next, we applied the two-way mixed ANOVA on the standard deviation of reproduced duration (i.e., precision), which revealed that neither Prior Task 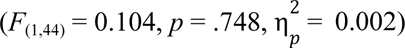, nor Cue Setting 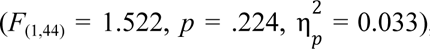, nor their interaction 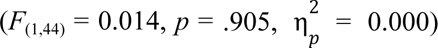 was significant. The results suggest that the time task difficulties for the two experiments were comparable.

**Figure 4.**
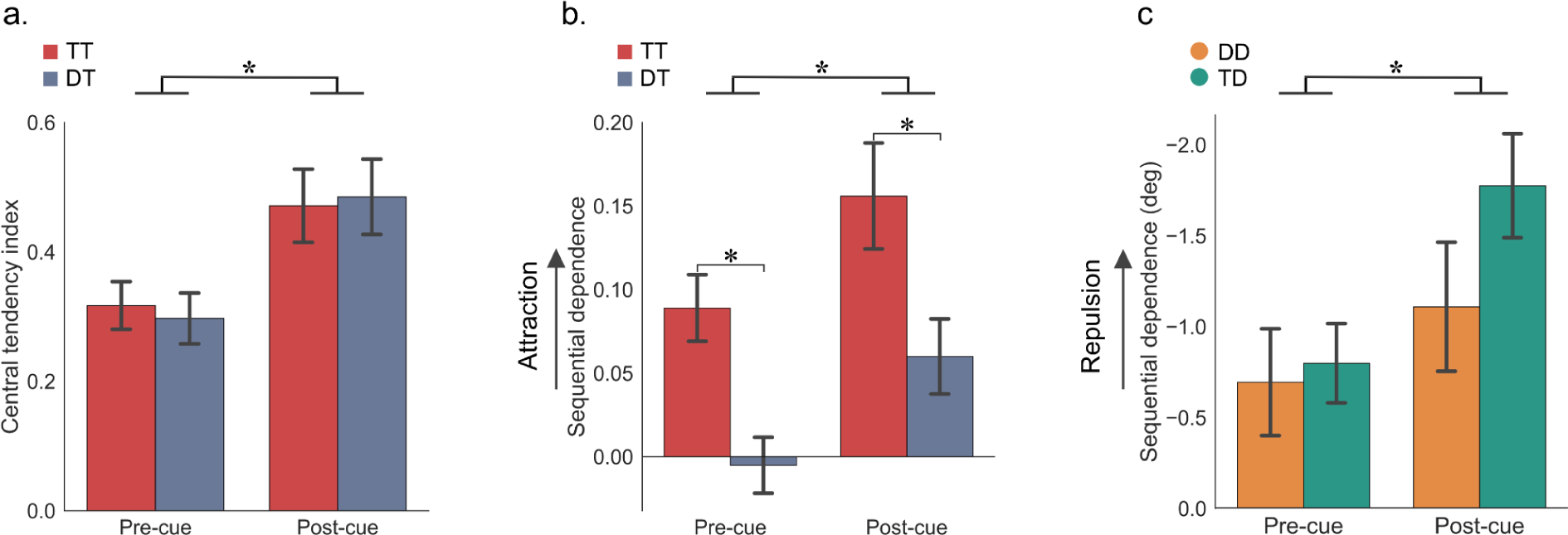
Comparisons between experiments (Exp. 1: pre-cue vs. Exp. 2: post-cue) and prior tasks (task-related vs. task-unrelated). (**a**) Central tendency effect for time reproduction trials. (**b**) Attractive sequential effect for time reproduction trials. TT represents consecutive time tasks, while DT represents a current time task preceded by a direction task. (**c**) Repulsive sequential effect for direction reproduction trials. DD represents consecutive direction tasks, while TD represents a direction task preceded by a time task. Error bars represent ± SEM. \**p* < .05.

For the direction reproduction, the two-way mixed ANOVA on the repulsion effect (averaged across 45.0° and 67.5°) revealed a significant main effect of Cue Setting 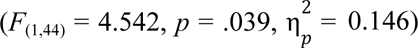. The repulsion effect was significantly enhanced with the post-cue relative to the pre-cue (Figure 4c). Neither Prior Task 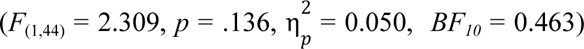 nor the interaction 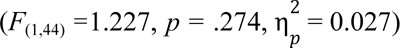 was significant. Another two-way mixed ANOVA for the attractive bias at an orientation difference of 90° revealed neither the main effect Prior Task nor Cue Setting, nor their interaction was significant (all *Fs* < 0.329, all *ps* > 0.569, 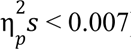). This means that the attractive bias at the orientation difference of 90° was not influenced by task relevance (task-relevant vs. task-irrelevant) or the cue position (pre-cueing vs. post-cueing).

In summary, comparison across experiments revealed that the post-cue condition enhanced the central tendency and sequential biases in time estimation, suggesting that the sequential attractive bias in time reproduction is influenced by working memory and post-perceptual processing, consistent with prior research (Bliss et al., 2017). In contrast, direction reproduction was unaffected by task relevance in both experiments, indicating that the repulsive bias likely originated from low-level sensory adaptation. The repulsion effects at 45.0° and 67.5° were enhanced by the post-cue (but not at the extreme case of 90°), possibly due to increased working memory load with the post-cue. Our findings contrast with those of Bae and Luck (2020), who reported differential effects of prior tasks with the post-cue setting. The discrepancy may arise from the fact that their experimental setup involved two visual tasks (color and direction) that might interact with each other in working memory, while we employed time and motion direction tasks.

## General Discussion

The present study investigated differential sequential effects in non-temporal and temporal tasks, using task changes and pre-cue vs. post-cue settings to examine the influences of task relevance and working memory load. Intriguingly, we observed only sequential attractive biases in timing tasks, but both repulsion and attraction effects in direction tasks across both experiments. For the time reproduction task, the attractive bias was reliable when the preceding trial involved the same task. However, this attraction was significantly reduced in the post-cue setting and vanished in the pre-cue setting when the preceding trial involved a direction task. In contrast, task relevance had no impact on sequential effects in direction reproduction. Nonetheless, the post-cue setting enhanced both attractive and repulsive biases.

Both sequential attractive and repulsive biases are well-documented in previous studies on orientation and direction judgments. For example, when stimuli such as Gabor orientations or gratings are used, small orientation differences between trials (under 20°) typically yield attraction biases, while larger differences elicit repulsion biases (Ceylan & Pascucci, 2023; Fritsche et al., 2017; Fritsche & de Lange, 2019a). Repulsive bias in orientation may originate from early sensory processing mechanisms, where neurons adapt to prolonged exposure to a specific orientation or direction, decreasing their sensitivity to that feature and subsequently shifting their spatial tuning, which causes negative tilt-aftereffect (Alais et al., 2017; Fritsche et al., 2020). Previous research has demonstrated that attractive and repulsive biases can occur concurrently, such as in motion direction processing (Alais et al., 2017; Ceylan & Pascucci, 2023; Fischer et al., 2020; Moon et al., 2022). For instance, a brief presentation (e.g., 200 ms in Alais et al., 2017 and Fischer et al., 2020) or mostly random motion display (Moon & Kwon, 2022) can make the orientation signal of the motion more dominant, resulting in attractive effects similar to those observed in static orientation studies (Fischer & Whitney, 2014; Manassi et al., 2018). In contrast, long exposure to a coherent motion signal (e.g., here 800 to 1600 ms) may induce motion adaptation, resulting in repulsive biases, similar to negative tilt-aftereffects (Alais et al., 2017; Moon et al., 2022). Our studies used coherent motion and found repulsion effects, consistent with recent studies on motion direction (Alais et al., 2017; Bae & Luck, 2017, 2020; Kang & Choi, 2015) or orientation (Ceylan & Pascucci, 2023; Su et al., 2023). Repulsive biases dominate when previous stimuli are either unattended or irrelevant to the task, or when visual stimuli have a long duration and high contrast, or a reference (Manassi et al., 2018; Pascucci et al., 2019; Pascucci & Plomp, 2021; Su et al., 2023). The repulsive bias observed here is likely due to dominant low-level motion adaptation with relatively long exposure times (800 to 1600 ms), which may overshadow any minor high-level task modulation, irrespective of the preceding duration or direction tasks.

The fact that repulsive biases can be enhanced by working memory load suggests that both early motion adaptation and late post-perceptual decision-making contribute to the observed sequential repulsion. The latter contribution indicates the involvement of high-level working memory processes, such as maintaining discriminability of multiple items (Czoschke et al., 2019; Fritsche et al., 2017), and active discarding of irrelevant information as well as reduced attention to irrelevant items (Ceylan & Pascucci, 2023). When memory load increases to reach the capacity limit, repulsive representation of multiple items could help maintain discriminability between items being held (Czoschke et al., 2019). This might also explain why repulsive biases occur away from unattended or irrelevant items (Ceylan & Pascucci, 2023; Fischer & Whitney, 2014; Shan & Postle, 2022). For example, irrelevant inducers, while initially attended to, are actively removed from working memory due to limited capacity (Lewis-Peacock et al., 2018; Shan & Postle, 2022). To protect the target item from removal, its representation is repelled away from those irrelevant inducers (Ceylan & Pascucci, 2023; Fritsche & de Lange, 2019b). In our study, the enhancement of the repulsive bias with the post-cue is likely due to the increased memory load, which pushes the current representation of the motion direction away from the preceding one. However, it should be noted that visual working memory load could increase the uncertainty of the target representation, which may result in an enhanced repulsion effect. The increased uncertainty may also potentially lead to an interaction of memory load and task modulation. For example, when the two tasks were in the same visual modality (i.e., orientation and color) and with the post-cue setting, the limited memory capacity may amplify the task relevance effect on repulsive biases (e.g., Bae & Luck, 2020).

Conversely, we observed attractive biases in the timing task. What accounts for the opposing patterns in sequential biases between timing and non-timing (direction) tasks? Time perception, unlike visual perception, lacks dedicated sensory systems (Wittmann & Paulus, 2008). The brain constructs time perception by integrating current sensory estimations with recent history and prior knowledge of the stimuli to enhance processing efficiency. This integration leads to attractive biases, as recent events serve as predictions (Glasauer & Shi, 2022; Shi et al., 2013). Glasauer and Shi (2022) showed that individual beliefs in temporal continuity impact the magnitudes of the sequential bias, with stronger attractive biases in those with high beliefs in temporal continuity. Unlike visual perception, which involves both early sensory adaptation and post-perceptual processing, time perception relies heavily on post-perceptual processes. In a duration reproduction task, reproducing duration relies on not only the encoded duration in working memory (Cheng et al., 2023) but also prior knowledge of the duration distribution (Jazayeri & Shadlen, 2010; Lejeune & Wearden, 2009; Shi & Burr, 2016). These sources are integrated and mixed in working memory to boost the reliability of estimates (Bausenhart et al., 2016; B. M. Gu & Meck, 2011; Penney et al., 2000). This memory mixing may involve active recall of memory traces from past experiences, contributing to sequential biases (Bliss et al., 2017; Ceylan & Pascucci, 2023; Fornaciai & Park, 2020; Fritsche & de Lange, 2019a; Ranieri et al., 2022).

For non-temporal visual processing, working memory may link sensory representation to responses, enhancing sequential effects. The sensory-level sequential effects can be either attractive or repulsive, depending on their functional roles (Ceylan et al., 2021; Fritsche & de Lange, 2019b; Glasauer & Shi, 2022; Kim et al., 2020; Kim & Alais, 2021; Suárez-Pinilla et al., 2018). For example, increasing the delay between stimulus presentation and response, thereby prolonging the reliance on working memory, leads to a stronger attractive bias toward the preceding stimulus (Bliss et al., 2017). On the other hand, when two visual orientations must be held in the working memory, their representations repel each other (Czoschke et al., 2019). In our study, increasing working memory load by using the post-cue enhanced both the adaptation-induced repulsive bias in direction judgments and attractive biases in time judgments, similar to previous findings that working memory can amplify sequential effects (Bliss et al., 2017).

One might ask about the exact underlying mechanism of this enhancement by working memory load. In our study, with the post-cue, both the non-temporal direction and the time interval had to be simultaneously encoded in working memory before the cue appeared. Shared memory representations likely increase sensory uncertainty (Li et al., 2021; Michail et al., 2021; Simon et al., 2016). According to Bayesian dynamic updating processes (Burr & Cicchini, 2014; Glasauer & Shi, 2022), the weight of prior stimuli increases during the integration, resulting in stronger sequential effects (Ceylan et al., 2021; Cicchini et al., 2018; Markov et al., 2024). Recent research also confirms that uncertainty can modulate the strength of serial dependence (Fulvio et al., 2023; Ozkirli & Pascucci, 2023). Further research is needed to clarify the exact mechanisms through which working memory load influences these sequential biases.

In summary, our study dissected sequential biases in space and time using a unified setting that tested both spatial motion direction and time reproduction. We uncovered distinct sequential biases: time blends through assimilation, while direction skews via dominant repulsion, with time particularly influenced by task relevance. Our findings highlight that sensory adaptation dominates repulsion biases in motion direction judgments, while post-perceptual processes that involve working memory have a greater effect on the bias in time reproduction. Moreover, increasing the working memory load with the post cue enhanced both opposing sequential effects. The distinct pattern of sequential biases between time and space potentially links to the different stages at which sequential effects emerge in processing non-temporal and temporal information.

## Data and code availability statement

The data and analysis code that support the findings of this study will be made available from the author, Si Cheng, upon reasonable request. All data and code will be made available in online repositories upon acceptance. This study was not preregistered.

## Acknowledgments

This study was supported by German Research Foundation (DFG) research grants SH 166/3-2 to Z. Shi and CH 3093/1-1 to S. Chen, and the Chinese CSC scholarship to S. Cheng.

## Footnotes

1 To enhance statistical power given the limited number of trials, we splitted durations into two categories: short (<1.2 s) and long (> 1.2 s).

## Reference

Alais, D., Leung, J., & Van der Burg, E. (2017). Linear Summation of Repulsive and Attractive Serial Dependencies: Orientation and Motion Dependencies Sum in Motion Perception. The Journal of Neuroscience: The Official Journal of the Society for Neuroscience, 37(16), 4381–4390.

Bae, G.-Y. (2024). Cardinal bias interacts with the stimulus history bias in orientation working memory. Attention, Perception & Psychophysics, 86(3), 828–837.

Bae, G.-Y., & Luck, S. J. (2017). Interactions between visual working memory representations. Attention, Perception & Psychophysics, 79(8), 2376–2395.

Bae, G.-Y., & Luck, S. J. (2019). Reactivation of Previous Experiences in a Working Memory Task. Psychological Science, 30(4), 587–595.

Bae, G.-Y., & Luck, S. J. (2020). Serial dependence in vision: Merely encoding the previous-trial target is not enough. Psychonomic Bulletin & Review, 27(2), 293–300.

Bausenhart, K. M., Bratzke, D., & Ulrich, R. (2016). Formation and representation of temporal reference information. Current Opinion in Behavioral Sciences, 8, 46–52.

Bliss, D. P., Sun, J. J., & D’Esposito, M. (2017). Serial dependence is absent at the time of perception but increases in visual working memory. Scientific Reports, 7(1), 1–13.

Burr, D. C., & Cicchini, G. M. (2014). Vision: Efficient Adaptive Coding. Current Biology: CB, 24(22), R1096–R1098.

Ceylan, G., Herzog, M. H., & Pascucci, D. (2021). Serial dependence does not originate from low-level visual processing. Cognition, 212, 104709.

Ceylan, G., & Pascucci, D. (2023). Attractive and repulsive serial dependence: The role of task relevance, the passage of time, and the number of stimuli. Journal of Vision, 23(6), 8.

Cheng, S., Chen, S., Glasauer, S., Keeser, D., & Shi, Z. (2023). Neural mechanisms of sequential dependence in time perception: the impact of prior task and memory processing. Cerebral Cortex. 10.1093/cercor/bhad453

Cicchini, G. M., Anobile, G., & Burr, D. C. (2014). Compressive mapping of number to space reflects dynamic encoding mechanisms, not static logarithmic transform. Proceedings of the National Academy of Sciences of the United States of America, 111(21), 7867–7872.

Cicchini, G. M., Arrighi, R., Cecchetti, L., Giusti, M., & Burr, D. C. (2012). Optimal encoding of interval timing in expert percussionists. The Journal of Neuroscience: The Official Journal of the Society for Neuroscience, 32(3), 1056–1060.

Cicchini, G. M., Mikellidou, K., & Burr, D. C. (2018). The functional role of serial dependence. Proceedings. Biological Sciences / The Royal Society, 285(1890). 10.1098/rspb.2018.1722

Cicchini, G. M., Mikellidou, K., & Burr, D. C. (2023). Serial Dependence in Perception. Annual Review of Psychology. 10.1146/annurev-psych-021523-104939

Czoschke, S., Fischer, C., Beitner, J., Kaiser, J., & Bledowski, C. (2019). Two types of serial dependence in visual working memory. British Journal of Psychology, 110(2), 256–267.

de Azevedo Neto, R. M., & Bartels, A. (2021). Disrupting Short-Term Memory Maintenance in Premotor Cortex Affects Serial Dependence in Visuomotor Integration. The Journal of Neuroscience: The Official Journal of the Society for Neuroscience, 41(45), 9392–9402.

Faul, F., Erdfelder, E., Lang, A.-G., & Buchner, A. (2007). G*Power 3: a flexible statistical power analysis program for the social, behavioral, and biomedical sciences. Behavior Research Methods, 39(2), 175–191.

Feigin, H., Baror, S., Bar, M., & Zaidel, A. (2021). Perceptual decisions are biased toward relevant prior choices. Scientific Reports, 11(1), 648.

Fischer, Czoschke, S., Peters, B., Rahm, B., Kaiser, J., & Bledowski, C. (2020). Context information supports serial dependence of multiple visual objects across memory episodes. Nature Communications, 11(1), 1932.

Fischer, & Whitney, D. (2014). Serial dependence in visual perception. Nature Neuroscience, 17(5), 738–743.

Fornaciai, M., & Park, J. (2018). Attractive Serial Dependence in the Absence of an Explicit Task. Psychological Science, 29(3), 437–446.

Fornaciai, M., & Park, J. (2019). Serial dependence generalizes across different stimulus formats, but not different sensory modalities. Vision Research, 160, 108–115.

Fornaciai, M., & Park, J. (2020). Neural Dynamics of Serial Dependence in Numerosity Perception. Journal of Cognitive Neuroscience, 32(1), 141–154.

Fritsche, M., & de Lange, F. P. (2019a). Reference repulsion is not a perceptual illusion. Cognition, 184, 107–118.

Fritsche, M., & de Lange, F. P. (2019b). The role of feature-based attention in visual serial dependence. Journal of Vision, 19(13), 21.

Fritsche, M., Mostert, P., & de Lange, F. P. (2017). Opposite Effects of Recent History on Perception and Decision. Current Biology: CB, 27(4), 590–595.

Fritsche, M., Spaak, E., & de Lange, F. P. (2020). A Bayesian and efficient observer model explains concurrent attractive and repulsive history biases in visual perception. eLife, 9, e55389.

Fulvio, J. M., Rokers, B., & Samaha, J. (2023). Task feedback suggests a post-perceptual component to serial dependence. Journal of Vision, 23(10), 6.

Girshick, A. R., Landy, M. S., & Simoncelli, E. P. (2011). Cardinal rules: visual orientation perception reflects knowledge of environmental statistics. Nature Neuroscience, 14(7), 926–932.

Glasauer, S., & Shi, Z. (2022). Individual beliefs about temporal continuity explain variation of perceptual biases. Scientific Reports, 12(1), 10746.

Gu, B. M., & Meck, W. H. (2011). New perspectives on Vierordt’s law: memory-mixing in ordinal temporal comparison tasks. In A. Vatakis, A. Esposito, M. Giagkou, F. Cummins, & G. Papadelis (Eds.), Multidisciplinary Aspects of Time and Time Perception (Vol. 6789, pp. 67–78).

Gu, B.-M., van Rijn, H., & Meck, W. H. (2015). Oscillatory multiplexing of neural population codes for interval timing and working memory. Neuroscience and Biobehavioral Reviews, 48, 160–185.

Hahn, M., & Wei, X.-X. (2024). A unifying theory explains seemingly contradictory biases in perceptual estimation. Nature Neuroscience. 10.1038/s41593-024-01574-x

Hayashi, M. J., Kantele, M., Walsh, V., Carlson, S., & Kanai, R. (2014). Dissociable neuroanatomical correlates of subsecond and suprasecond time perception. Journal of Cognitive Neuroscience, 26(8), 1685–1693.

Holland, M. K., & Lockhead, G. R. (1968). Sequential effects in absolute judgments of loudness. Perception & Psychophysics, 3(6), 409–414.

Jazayeri, M., & Shadlen, M. N. (2010). Temporal context calibrates interval timing. Nature Neuroscience, 13(8), 1020–1026.

Jesteadt, W., Luce, R. D., & Green, D. M. (1977). Sequential effects in judgments of loudness. Journal of Experimental Psychology. Human Perception and Performance, 3(1), 92–104.

Kang, M.-S., & Choi, J. (2015). Retrieval-Induced Inhibition in Short-Term Memory. Psychological Science, 26(7), 1014–1025.

Kim, S., & Alais, D. (2021). Individual differences in serial dependence manifest when sensory uncertainty is high. Vision Research, 188, 274–282.

Kim, S., Burr, D., Cicchini, G. M., & Alais, D. (2020). Serial dependence in perception requires conscious awareness. Current Biology: CB, 30(6), R257–R258.

Kiyonaga, A., Manassi, M., & Whitney, D. (2017). Context transitions modulate perceptual serial dependence. Journal of Vision, 17(10), 92–92.

Kiyonaga, A., Scimeca, J. M., Bliss, D. P., & Whitney, D. (2017). Serial Dependence across Perception, Attention, and Memory. Trends in Cognitive Sciences, 21(7), 493–497.

Lejeune, H., & Wearden, J. H. (2009). Vierordt’sThe Experimental Study of the Time Sense(1868) and its legacy. The European Journal of Cognitive Psychology, 21(6), 941–960.

Lewis, P. A., & Miall, R. C. (2003). Brain activation patterns during measurement of sub-and supra-second intervals. Neuropsychologia, 41(12), 1583–1592.

Lewis-Peacock, J. A., Kessler, Y., & Oberauer, K. (2018). The removal of information from working memory. Annals of the New York Academy of Sciences, 1424(1), 33–44.

Liberman, A., Zhang, K., & Whitney, D. (2016). Serial dependence promotes object stability during occlusion. Journal of Vision, 16(15), 16.

Li, H.-H., Sprague, T. C., Yoo, A. H., Ma, W. J., & Curtis, C. E. (2021). Joint representation of working memory and uncertainty in human cortex. Neuron, 109(22), 3699–3712.e6.

Manassi, M., Liberman, A., Kosovicheva, A., Zhang, K., & Whitney, D. (2018). Serial dependence in position occurs at the time of perception. Psychonomic Bulletin & Review, 25(6), 2245–2253.

Manassi, M., Murai, Y., & Whitney, D. (2023). Serial dependence in visual perception: A meta-analysis and review. Journal of Vision, 23(8), 18.

Mao, J., & Stocker, A. (2021). Orientation perception is based on efficient coding and categorical decoding. Journal of Vision, 21(9), 2643–2643.

Markov, Y. A., Tiurina, N. A., & Pascucci, D. (2024). Serial dependence: A matter of memory load. Heliyon, 10(13), e33977.

Michail, G., Senkowski, D., Niedeggen, M., & Keil, J. (2021). Memory Load Alters Perception-Related Neural Oscillations during Multisensory Integration. The Journal of Neuroscience: The Official Journal of the Society for Neuroscience, 41(7), 1505–1515.

Moon, J., & Kwon, O.-S. (2022). Attractive and repulsive effects of sensory history concurrently shape visual perception. BMC Biology, 20(1), 247.

Moon, J., Tadin, D., & Kwon, O.-S. (2022). A key role of orientation in the coding of visual motion direction. Psychonomic Bulletin & Review. 10.3758/s13423-022-02181-2

Murai, Y., & Whitney, D. (2021). Serial dependence revealed in history-dependent perceptual templates. Current Biology: CB, 31(14), 3185–3191.e3.

Ozkirli, A., & Pascucci, D. (2023). State-dependent serial dependence in perceptual decisions. In bioRxiv (p. 2023.10.19.563128). 10.1101/2023.10.19.563128

Pascucci, D., Mancuso, G., Santandrea, E., Della Libera, C., Plomp, G., & Chelazzi, L. (2019). Laws of concatenated perception: Vision goes for novelty, decisions for perseverance. PLoS Biology, 17(3), e3000144.

Pascucci, D., & Plomp, G. (2021). Serial dependence and representational momentum in single-trial perceptual decisions. Scientific Reports, 11(1), 9910.

Pascucci, D., Roinishvili, M., Chkonia, E., Brand, A., Whitney, D., Herzog, M. H., & Manassi, M. (2024). Intact Serial Dependence in Schizophrenia: Evidence from an Orientation Adjustment Task. Schizophrenia Bulletin. 10.1093/schbul/sbae106

Pascucci, D., Tanrikulu, Ö. D., Ozkirli, A., Houborg, C., Ceylan, G., Zerr, P., Rafiei, M., & Kristjánsson, Á. (2023). Serial dependence in visual perception: A review. Journal of Vision, 23(1), 9.

Peirce, J., Gray, J. R., Simpson, S., MacAskill, M., Höchenberger, R., Sogo, H., Kastman, E., & Lindeløv, J. K. (2019). PsychoPy2: Experiments in behavior made easy. Behavior Research Methods, 51(1), 195–203.

Penney, T. B., Gibbon, J., & Meck, W. H. (2000). Differential Effects of Auditory and Visual Signals on Clock Speed and Temporal Memory. Journal of Experimental Psychology. Human Perception and Performance, 26(6), 1770.

Ranieri, G., Benedetto, A., Ho, H. T., Burr, D. C., & Morrone, M. C. (2022). Evidence of Serial Dependence from Decoding of Visual Evoked Potentials. The Journal of Neuroscience: The Official Journal of the Society for Neuroscience, 42(47), 8817–8825.

Sadil, P., Cowell, R. A., & Huber, D. E. (2024). The push-pull of serial dependence effects: Attraction to the prior response and repulsion from the prior stimulus. Psychonomic Bulletin & Review, 31(1), 259–273.

Samaha, J., Switzky, M., & Postle, B. R. (2019). Confidence boosts serial dependence in orientation estimation. Journal of Vision, 19(4), 25. https://osf.io/6uczk/.

Shan, J., & Postle, B. R. (2022). The Influence of Active Removal from Working Memory on Serial Dependence. Journal of Cognition, 5(1), 31.

Sheehan, T. C., & Serences, J. T. (2022). Attractive serial dependence overcomes repulsive neuronal adaptation. PLoS Biology, 20(9), e3001711.

Sheehan, T. C., & Serences, J. T. (2023). Distinguishing response from stimulus driven history biases. In bioRxiv (p. 2023.01.11.523637). 10.1101/2023.01.11.523637

Shi, Z., & Burr, D. (2016). Predictive coding of multisensory timing. Current Opinion in Behavioral Sciences, 8, 200–206.

Shi, Z., Church, R. M., & Meck, W. H. (2013). Bayesian optimization of time perception. Trends in Cognitive Sciences, 17(11), 556–564.

Simon, S. S., Tusch, E. S., Holcomb, P. J., & Daffner, K. R. (2016). Increasing Working Memory Load Reduces Processing of Cross-Modal Task-Irrelevant Stimuli Even after Controlling for Task Difficulty and Executive Capacity. Frontiers in Human Neuroscience, 10, 380.

Suárez-Pinilla, M., Seth, A. K., & Roseboom, W. (2018). Serial dependence in the perception of visual variance. Journal of Vision, 18(7), 4.

Su, Y., Wachtler, T., & Shi, Z. (2023). Reference induces biases in late visual processing. Scientific Reports, 13(1), 18624.

Teki, S., & Griffiths, T. D. (2016). Brain Bases of Working Memory for Time Intervals in Rhythmic Sequences. Frontiers in Neuroscience, 10, 239.

Wehrman, Sanders, R., & Wearden, J. (2023). What came before: Assimilation effects in the categorization of time intervals. Cognition, 234, 105378.

Wehrman, Wearden, J. H., & Sowman, P. (2018). Short-term effects on temporal judgement: Sequential drivers of interval bisection and reproduction. Acta Psychologica, 185, 87–95.

Wiener, M., Thompson, J. C., & Coslett, H. B. (2014). Continuous carryover of temporal context dissociates response bias from perceptual influence for duration. PloS One, 9(6), e100803.

Wittmann, M., & Paulus, M. P. (2008). Decision making, impulsivity and time perception. Trends in Cognitive Sciences, 12(1), 7–12.

Zhou, L., Liu, Y., Jiang, Y., Wang, W., Xu, P., & Zhou, K. (2024). The distinct development of stimulus and response serial dependence. Psychonomic Bulletin & Review. 10.3758/s13423-024-02474-8

